# RstA Regulation of *Clostridioides difficile* Toxin Production and Sporulation in Phenotypically Diverse Strains

**DOI:** 10.1101/774125

**Authors:** Adrianne N. Edwards, Ellen G. Krall, Shonna M. McBride

## Abstract

The anaerobic spore-former, *Clostridioides difficile*, causes significant diarrheal disease in humans and other mammals. Infection begins with the ingestion of dormant spores which subsequently germinate within the host gastrointestinal tract. Here, the vegetative cells proliferate and secrete two exotoxins, TcdA and TcdB, which cause disease symptoms. Although spore formation and toxin production are critical for *C. difficile* pathogenesis, the regulatory links between these two physiological processes are not well understood and are strain-dependent. Previously, we identified a conserved *C. difficile* regulator, RstA, that promotes sporulation initiation through an unknown mechanism and directly and indirectly represses toxin gene transcription in the historical isolate, 630Δ*erm*. To test whether perceived strain-dependent differences in toxin production and sporulation are mediated by RstA, we created an *rstA* mutant in the epidemic 027 ribotype strain, R20291. RstA affects sporulation and toxin gene expression similarly in R20291, although more robust regulatory effects are observed in this strain than in 630Δ*erm*. Reporter assays measuring transcriptional regulation of *tcdR*, the sigma factor essential for toxin gene expression, identified sequence-dependent effects influencing repression by RstA and CodY, a global nutritional sensor, in four diverse *C. difficile* strains. We provide evidence that RstA contributes to *tcdR* bistability in R20291 by biasing cells to a toxin-OFF state. Finally, sequence-dependent and strain-dependent differences were evident in RstA negative autoregulation of *rstA* transcription. Our data establish RstA as an important regulator of *C. difficile* virulence traits, and implicate RstA as a contributor to the variety of sporulation and toxin phenotypes observed in distinct isolates.

**IMPORTANCE:** Two critical traits of *Clostridioides difficile* pathogenesis are the production of toxins, which cause disease symptoms, and the formation of spores, which permit survival outside of the gastrointestinal tract. The multifunctional regulator, RstA, promotes sporulation and prevents toxin production in the historical strain, 630Δ*erm*. Here, we show that RstA functions similarly in an epidemic isolate, R20291, although strain-specific effects on toxin and *rstA* expression are evident. Our data demonstrate that sequence-specific differences within the promoter for the toxin regulator, TcdR, contribute to regulation of toxin production by RstA and CodY. These sequence differences account for some of the variability in toxin production among isolates, and may allow strains to differentially control toxin production in response to a variety of signals.

## INTRODUCTION

As the leading cause of antibiotic-associated diarrhea, *Clostridioides difficile* resides in the mammalian gastrointestinal tract, where disease symptoms are mediated by the production of two large, glucosylating exotoxins, toxin A (TcdA) and toxin B (TcdB) (1). TcdA and TcdB target the Rho and Ras family of small GTPases (2, 3), ultimately disrupting host cell function and triggering apoptotic and/or necrotic cell death (4). TcdA and TcdB are encoded within the 19.6 kb pathogenicity locus (PaLoc) along with *tcdR*, the toxin-specific sigma factor that is required for toxin gene expression, *tcdE*, encoding a holin that likely allows for toxin efflux, and *tcdC*, a putative negative regulator of toxin production whose function is debated (5-10). Because synthesis of these large toxins is energy-intensive, *C. difficile* toxin gene expression is directly repressed by multiple regulatory factors to ensure that toxin production occurs only under conditions in which the function of the toxins contribute to the survival of the bacterium within the host (11-13).

Additionally, as a strict anaerobe, *C. difficile* relies on the morphological transformation into a dormant spore to survive the subsequent exodus from the gastrointestinal tract and efficient transmission to a new host (14). While the characteristic morphological stages of sporulation are conserved, the regulatory network that controls *C. difficile* sporulation initiation, and thus, the activation of Spo0A, the master regulator of sporulation, is divergent from other spore formers and is poorly mapped out (15). The three transcriptional repressors, CodY, CcpA and RstA, that directly repress toxin gene expression in *C. difficile*, are also involved in the regulation of early stage sporulation events; however, the genetic pathways through which these regulators control spore formation have not been fully delineated (16-18). CodY and CcpA are largely transcriptional repressors that coordinate adaptation to environmental conditions in response to nutritional signals (17, 19). RstA is a member of the multifunctional RRNPP family of proteins and controls toxin production and sporulation through separate domains (13, 18), indicating that RstA utilizes two different molecular mechanisms to regulate these phenotypes. RstA function has only been studied in the laboratory isolate, 630Δ*erm*, and its contribution to sporulation and toxin production has not been evaluated in other *C. difficile* strains.

As new *C. difficile* PCR ribotypes emerge and prevail in the clinical population, the toxin and sporulation phenotypes of these isolates are often characterized to determine which traits allow these strains to exhibit increased virulence and circulate persistently (20-25). The variability in *tcdA* and *tcdB* gene sequences has led to the established method of toxinotyping *C. difficile* strains using PCR-restriction fragment length polymorphisms (RFLPs; reviewed in 26), although single nucleotide polymorphisms (SNPs) and small insertions and deletions located within the promoter regions and open reading frames of *tcdR, tcdE* and *tcdC* also purportedly contribute to toxin gene expression, production and secretion. Some of these changes have been documented in the literature, including deletions and frameshift mutations within the *tcdC* putative negative regulator (27, 28) and alternate TcdE isoforms that influence toxin secretion (29). Although there are a few nucleotide changes among *C. difficile* strains within the *tcdA* and *tcdB* promoter regions, none of these overlap the σ^TcdR^-dependent promoters essential for their transcription. However, numerous point mutations are located within the *tcdR* promoter region, many of which overlap the consensus sequences of the σ^A^- and σ^D^-dependent promoters and the RstA and CodY binding sites. We hypothesized that the point mutations within the *tcdR* promoters affect transcription initiation and influence RstA- and CodY-dependent repression, both of which may account for some of the variable, strain-specific toxin expression phenotypes observed.

To determine the impact RstA exerts on sporulation and toxin production in clinically-relevant *C. difficile* strains, a null *rstA* mutant was created in R20291, an epidemic isolate that emerged in the mid-2000s (30). We demonstrate that RstA is a regulator of critical virulence factors in this epidemic background and reveal strain-dependent differences that result in robust regulation of R20291 sporulation and toxin production. Further, we dissect the conserved regulatory and promoter features that control transcription of the bistable toxin regulator, *tcdR*, and define the regulatory contributions of RstA and CodY on toxin production in four important *C. difficile* strains. The observation that strain-specific nucleotide substitutions in the *tcdR* promoter alter regulation of toxin gene expression *in vitro* illuminates another molecular mechanism by which more virulent strains exhibit altered toxin levels during *C. difficile* infection.

## MATERIALS AND METHODS

### Bacterial strains and growth conditions

The bacterial strains and plasmids used in this study are found in **Table 1**. *C. difficile* strains were cultured in BHIS (31) or TY medium, pH 7.4 (32), in the presence of 2-5 *µ*g/ml thiamphenicol or 2-5 *µ*g/ml erythromycin, as indicated, in a 37°C anaerobic chamber containing an atmosphere of 10% H_2_, 5% CO_2_ and 85% N_2_, as previously detailed (33). *C. difficile* overnight cultures contained 0.1% taurocholate, to germinate spores, and 0.2% fructose, to prevent the formation of spores (32, 34). *Escherichia coli* and *Bacillus subtilis* strains were grown in LB (35), supplemented with 100 *µ*g/ml ampicillin and/or 20 *µ*g/ml chloramphenicol as indicated for *E. coli* or with 1 *µ*g/ml erythromycin, as indicated for *B. subtilis* at 37°C. After conjugation with *C. difficile, E. coli* and *B. subtilis* were counterselected using 50-100 *µ*g/ml kanamycin (36).

**Table 1.**
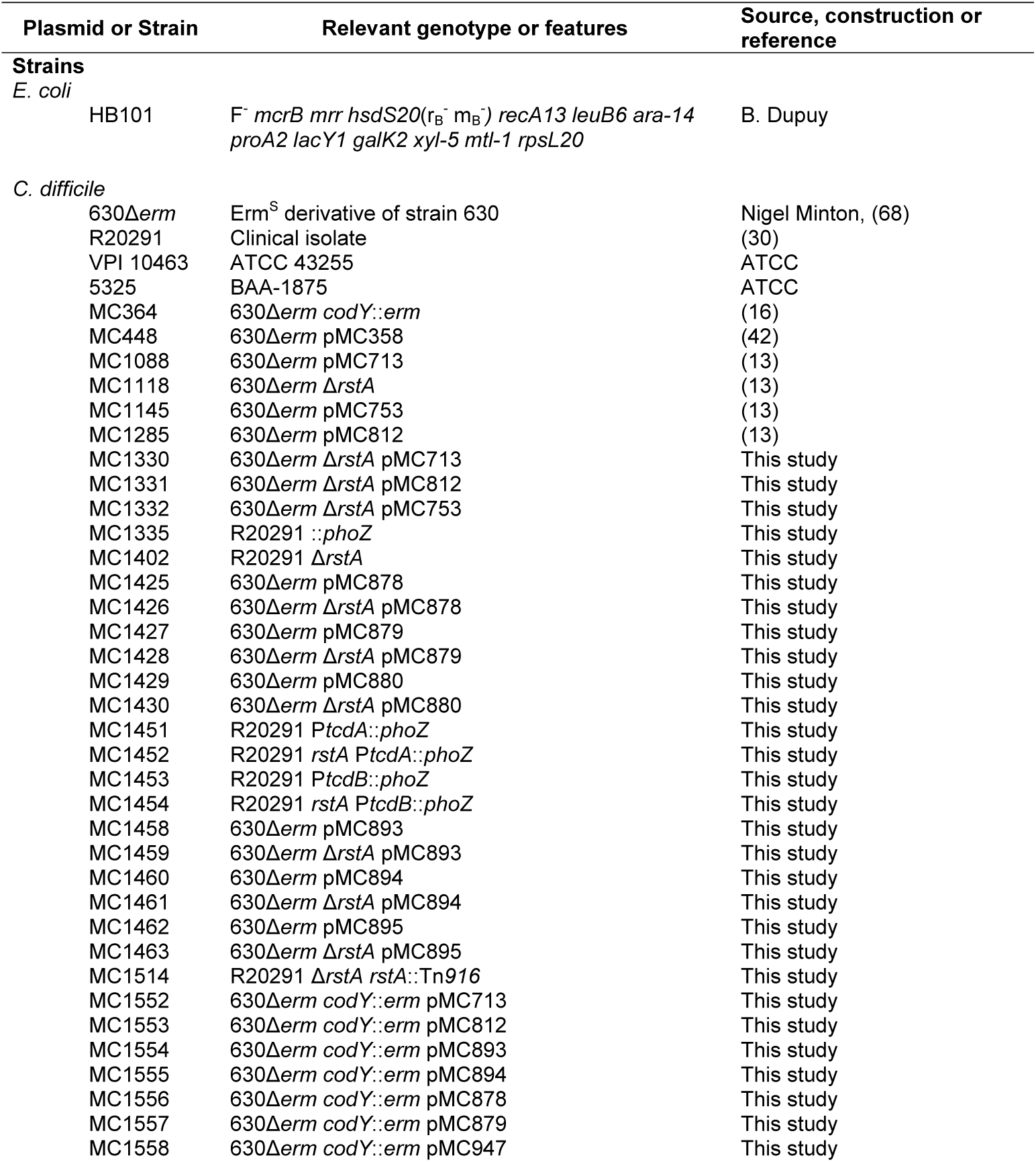
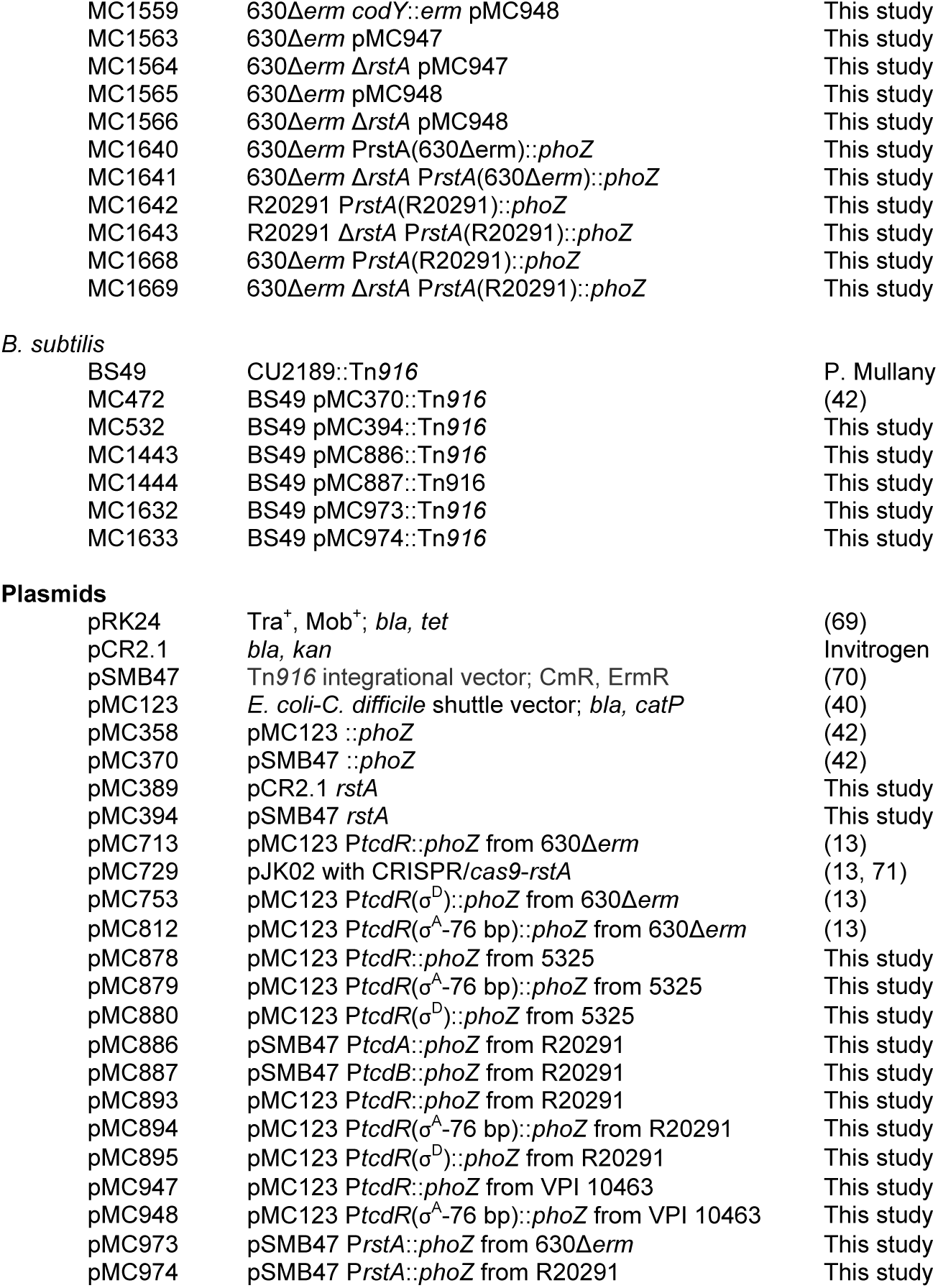
Bacterial Strains and plasmids.

| Plasmid or Strain | Relevant genotype or features | Source, construction or reference |
| --- | --- | --- |
| **Strains** |  |  |
| *E. coli* |  |  |
| HB101 | F<sup>-</sup> *mcrB mrr hsdS20(r<sub>B</sub><sup>-</sup> m<sub>B</sub><sup>-</sup>) recA13 leuB6 ara-14 proA2 lacY1 galK2 xyl-5 mtl-1 rpsL20* | B. Dupuy |
| *C. difficile* |  |  |
| 630Δ*erm* | Erm<sup>S</sup> derivative of strain 630 | Nigel Minton, (68) |
| R20291 | Clinical isolate | (30) |
| VPI 10463 | ATCC 43255 | ATCC |
| 5325 | BAA-1875 | ATCC |
| MC364 | 630Δ*erm* *codY::erm* | (16) |
| MC448 | 630Δ*erm* pMC358 | (42) |
| MC1088 | 630Δ*erm* pMC713 | (13) |
| MC1118 | 630Δ*erm* Δ*rstA* | (13) |
| MC1145 | 630Δ*erm* pMC753 | (13) |
| MC1285 | 630Δ*erm* pMC812 | (13) |
| MC1330 | 630Δ*erm* Δ*rstA* pMC713 | This study |
| MC1331 | 630Δ*erm* Δ*rstA* pMC812 | This study |
| MC1332 | 630Δ*erm* Δ*rstA* pMC753 | This study |
| MC1335 | R20291 ::*phoZ* | This study |
| MC1402 | R20291 Δ*rstA* | This study |
| MC1425 | 630Δ*erm* pMC878 | This study |
| MC1426 | 630Δ*erm* Δ*rstA* pMC878 | This study |
| MC1427 | 630Δ*erm* pMC879 | This study |
| MC1428 | 630Δ*erm* Δ*rstA* pMC879 | This study |
| MC1429 | 630Δ*erm* pMC880 | This study |
| MC1430 | 630Δ*erm* Δ*rstA* pMC880 | This study |
| MC1451 | R20291 *PtcdA::phoZ* | This study |
| MC1452 | R20291 *rstA PtcdA::phoZ* | This study |
| MC1453 | R20291 *PtcdB::phoZ* | This study |
| MC1454 | R20291 *rstA PtcdB::phoZ* | This study |
| MC1458 | 630Δ*erm* pMC893 | This study |
| MC1459 | 630Δ*erm* Δ*rstA* pMC893 | This study |
| MC1460 | 630Δ*erm* pMC894 | This study |
| MC1461 | 630Δ*erm* Δ*rstA* pMC894 | This study |
| MC1462 | 630Δ*erm* pMC895 | This study |
| MC1463 | 630Δ*erm* Δ*rstA* pMC895 | This study |
| MC1514 | R20291 Δ*rstA* *rstA::Tn916* | This study |
| MC1552 | 630Δ*erm* *codY::erm* pMC713 | This study |
| MC1553 | 630Δ*erm* *codY::erm* pMC812 | This study |
| MC1554 | 630Δ*erm* *codY::erm* pMC893 | This study |
| MC1555 | 630Δ*erm* *codY::erm* pMC894 | This study |
| MC1556 | 630Δ*erm* *codY::erm* pMC878 | This study |
| MC1557 | 630Δ*erm* *codY::erm* pMC879 | This study |
| MC1558 | 630Δ*erm* *codY::erm* pMC947 | This study |
| MC1559 | 630Δerm *codY::erm* pMC948 | This study |
| MC1563 | 630Δerm pMC947 | This study |
| MC1564 | 630Δerm ΔrstA pMC947 | This study |
| MC1565 | 630Δerm pMC948 | This study |
| MC1566 | 630Δerm ΔrstA pMC948 | This study |
| MC1640 | 630Δerm PrstA(630Δerm)::*phoZ* | This study |
| MC1641 | 630Δerm ΔrstA PrstA(630Δerm)::*phoZ* | This study |
| MC1642 | R20291 PrstA(R20291)::*phoZ* | This study |
| MC1643 | R20291 ΔrstA PrstA(R20291)::*phoZ* | This study |
| MC1668 | 630Δerm PrstA(R20291)::*phoZ* | This study |
| MC1669 | 630Δerm ΔrstA PrstA(R20291)::*phoZ* | This study |
| *B. subtilis* |  |  |
| BS49 | CU2189::Tn916 | P. Mullany |
| MC472 | BS49 pMC370::Tn916 | (42) |
| MC532 | BS49 pMC394::Tn916 | This study |
| MC1443 | BS49 pMC886::Tn916 | This study |
| MC1444 | BS49 pMC887::Tn916 | This study |
| MC1632 | BS49 pMC973::Tn916 | This study |
| MC1633 | BS49 pMC974::Tn916 | This study |
| Plasmids |  |  |
| pRK24 | Tra<sup>+</sup>, Mob<sup>+</sup>; *bla*, *tet* | (69) |
| pCR2.1 | *bla*, *kan* | Invitrogen |
| pSMB47 | Tn916 integrational vector; CmR, ErmR | (70) |
| pMC123 | *E. coli*-*C. difficile* shuttle vector; *bla*, *catP* | (40) |
| pMC358 | pMC123 ::*phoZ* | (42) |
| pMC370 | pSMB47 ::*phoZ* | (42) |
| pMC389 | pCR2.1 *rstA* | This study |
| pMC394 | pSMB47 *rstA* | This study |
| pMC713 | pMC123 *PtcdR*::*phoZ* from 630Δerm | (13) |
| pMC729 | pJK02 with CRISPR/*cas9-rstA* | (13, 71) |
| pMC753 | pMC123 *PtcdR*($\sigma^D$)::*phoZ* from 630Δerm | (13) |
| pMC812 | pMC123 *PtcdR*($\sigma^A$-76 bp)::*phoZ* from 630Δerm | (13) |
| pMC878 | pMC123 *PtcdR*::*phoZ* from 5325 | This study |
| pMC879 | pMC123 *PtcdR*($\sigma^A$-76 bp)::*phoZ* from 5325 | This study |
| pMC880 | pMC123 *PtcdR*($\sigma^D$)::*phoZ* from 5325 | This study |
| pMC886 | pSMB47 *PtcdA*::*phoZ* from R20291 | This study |
| pMC887 | pSMB47 *PtcdB*::*phoZ* from R20291 | This study |
| pMC893 | pMC123 *PtcdR*::*phoZ* from R20291 | This study |
| pMC894 | pMC123 *PtcdR*($\sigma^A$-76 bp)::*phoZ* from R20291 | This study |
| pMC895 | pMC123 *PtcdR*($\sigma^D$)::*phoZ* from R20291 | This study |
| pMC947 | pMC123 *PtcdR*::*phoZ* from VPI 10463 | This study |
| pMC948 | pMC123 *PtcdR*($\sigma^A$-76 bp)::*phoZ* from VPI 10463 | This study |
| pMC973 | pSMB47 PrstA::*phoZ* from 630Δerm | This study |
| pMC974 | pSMB47 PrstA::*phoZ* from R20291 | This study |

### Strain and plasmid construction

*C. difficile* strains 630 (RT012; Genbank no. NC_009089.1), R20291 (RT027; Genbank no. FN545816.1), VPI 10643 (ATCC 43255; RT078; Genbank no. NZ_CM000604.1) and M120 (RT078; Genbank no. NZ_FRES01000002.1) were used as templates for primer design and for PCR amplification, with the exception that 5325 (ATCC 1875; RT078; unsequenced) genomic DNA was the template for PCR amplification. Sequences of cloned PCR fragments were performed by Eurofins Genomics (Louisville, KY), and all M120 and 5325 sequences used were identical. Sequencing of the *C. difficile tcdE* open reading frames was performed by Genscript (Piscataway, NJ). Oligonucleotides used for PCR and qRT-PCR analyses are listed in **Table 2**. Details of strain and plasmid construction are detailed in the Supplemental Material (**Fig. S1**).

**Table 2.**
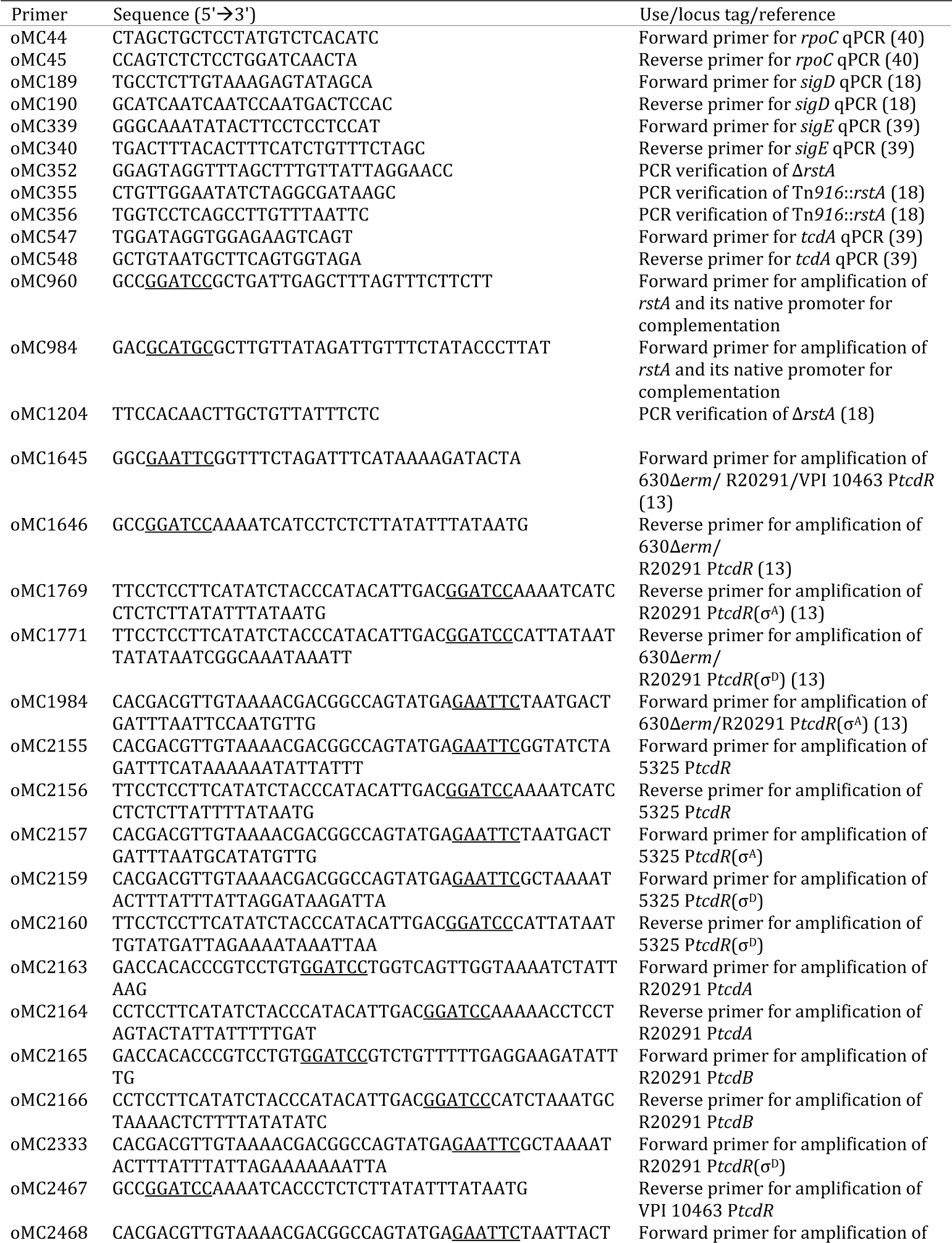
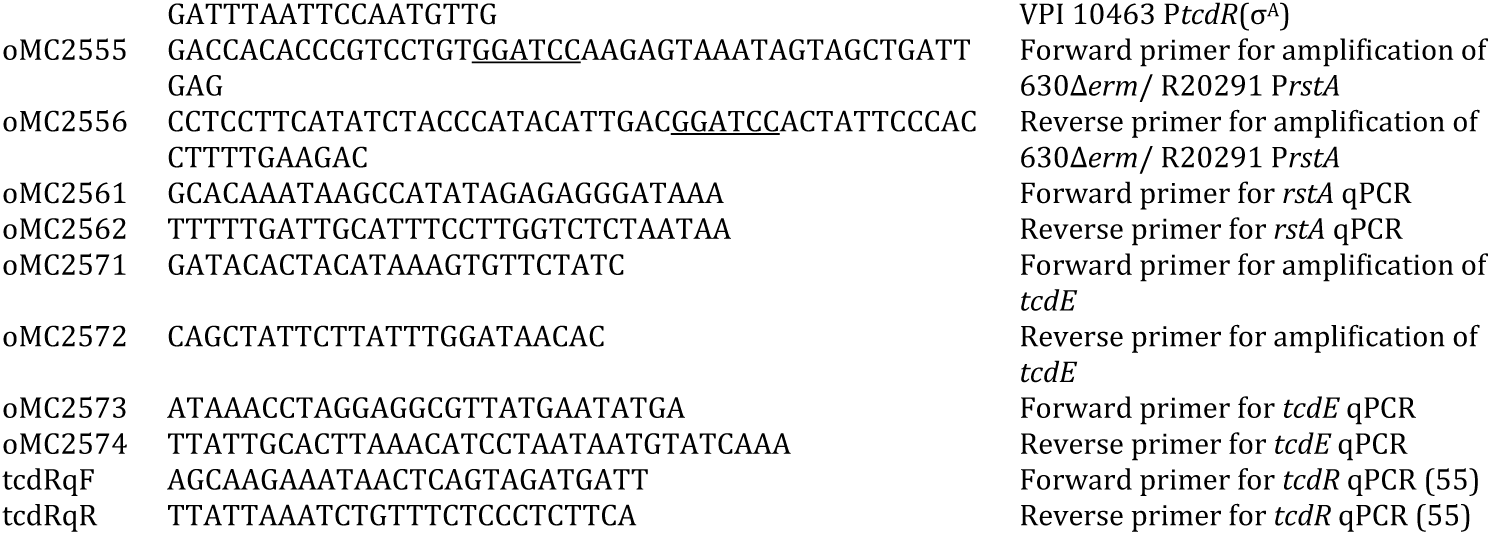
Oligonucleotides. All primers were acquired from IDT (Coralville, IA). Restriction enzyme sequences are underlined.

### Sporulation assays

Sporulation assays were performed as previously documented (37, 38). Briefly, *C. difficile* were cultured in BHIS medium supplemented with 0.1% taurocholate with 0.2% fructose until mid-exponential phase, and 0.25 ml culture was plated onto fresh 70:30 sporulation agar. After 24 h, the cells were removed from the plate and suspended in BHIS medium to an OD_600_ of ∼1.0. The total number of vegetative cells were determined by immediately plating serial dilutions of the suspended cells onto BHIS plates. To enumerate the total number of spores at the same time, an aliquot of suspended cells was mixed and incubated for 15 min with 28.5% ethanol to eliminate all vegetative cells and subsequently serially diluted in 1X PBS + 0.1% taurocholate and plated onto BHIS supplemented with 0.1% taurocholate. CFU were enumerated after at least 24 h of growth, and the sporulation frequency was calculated as the total number of ethanol-resistant spores divided by the total number of viable cells (spores plus vegetative cells). A *spo0A* mutant (MC310) was used as a negative sporulation control. Statistical significance was determined using a two-tailed Student’s *t-*test comparing each mutant to its isogenic parent.

### Phase contrast microscopy

At the same time cells were harvested for sporulation assays, as described above, an aliquot was removed from the anaerobic chamber, pelleted and suspended in ∼10 μl of supernatant. As detailed previously (39), the concentration culture was applied to a 0.7% agarose pad on the surface of the slide and imaged using a x100 Ph3 oil immersion objective on a Nikon Eclipse Ci-L microscope with a DS-Fi2 camera.

### Quantitative reverse transcription PCR analysis (qRT-PCR)

*C. difficile* was grown in either TY medium, pH 7.4 or on 70:30 sporulation agar, as described above. Cells were harvested from TY at T4, defined as four hours after the entry into stationary phase, or from 70:30 sporulation agar at H_12_, defined as 12 hours after spotting onto the plates. Cells harvested from TY were mixed with either 3 ml ice cold ethanol:acetone (1:1), or cells were scraped from plates and suspended in 6 ml ice-cold water:ethanol:acetone (3:1.5:1.5) and stored at -80°C. RNA was purified and DNase-I treated (Ambion) as previously detailed (19, 39, 40), and cDNA was synthesized using random hexamers (Bioline; 39). Quantitative reverse-transcription PCR (qRT-PCR) was performed in technical triplicate with 50 ng cDNA, using the SensiFAST SYBR & Fluorescein Kit (Bioline), on a Roche Lightcycler 96 for four biological replicates. To confirm the absence of contaminating genomic DNA, cDNA synthesis reactions containing no reverse transcriptase were included for all samples. Results were calculated using the comparative cycle threshold method (41), normalizing expression to the internal control transcript, *rpoC*. Either the two-tailed Student’s *t*-test was used to compare the activity in the *rstA* mutant to the parent strain or a one-way ANOVA followed by Tukey’s multiple comparisons test used, as indicated in the figure legends.

### Western blot analysis

Briefly, *C. difficile* strains were grown in TY medium, pH 7.4, and harvested at H_24_, defined as 24 hours after inoculation (13). Lysates were prepared and total protein quantitated using the Pierce Micro BCA Protein Assay Kit, as previously described (37). Total protein (3 *µ*g) was separated by electrophoresis on a precast 4-15% TGX Stain-free gradient gel (Bio-Rad), imaged using a ChemiDoc (Bio-Rad), and transferred to a 0.45 *µ*m nitrocellulose membrane. Western blot analysis was conducted with mouse anti-TcdA (Novus Biologicals) followed by goat anti-mouse Alexa Fluor 488 (Life Technologies) secondary antibody. Imaging and densitometry were performed with a ChemiDoc and Image Lab Software (Bio-Rad), and analyzed using a two-tailed Student’s *t-*test, comparing each mutant to its isogenic parent. A minimum of three biological replicates were analyzed for each strain, and a representative western blot image is shown.

### Enzyme-linked immunosorbent assay (ELISA)

To quantify *C. difficile* TcdA and TcdB in the supernatants of *C. difficile* cultures, aliquots of *C. difficile* strains grown in TY medium, pH 7.4, were collected at H_24_, pelleted, and supernatants diluted in provided Dilution Buffer were assayed using the Simultaneous Detection of *C. difficile* toxins A and B kit from TGCbiomics (TGC-E001-1), per manufacturer’s instructions. Technical duplicates were averaged and normalized to the OD_600_ of the respective cultures at H_24_, and the results are provided as the mean and standard error of the mean of three biological replicates. Statistical significance was determined using a two-tailed Student’s *t-*test, comparing each mutant to its isogenic parent.

### Alkaline phosphatase (AP) activity assays

Briefly, *C. difficile* strains were grown in TY medium, pH 7.4, and harvested at H_24_ for the P*tcdR* fusions or grown on 70:30 sporulation agar and harvested at H_8_ for the P*rstA* fusions (13). AP assays were performed as previously detailed (42), in the absence of chloroform. Technical duplicates were averaged, and the results are provided as the mean and standard error of the mean of at least three biological replicates. Either the two-tailed Student’s *t*-test was used to compare the activity in the *rstA* mutant to the parent strain or one-way ANOVA followed by Dunnett’s or Tukey’s multiple comparisons tests were performed, as indicated in the figure legends.

## RESULTS

### RstA positively influences R20291 sporulation

To assess the contribution of RstA in controlling sporulation and toxin production in phenotypically diverse *C. difficile* strains, we attempted to create null *rstA* mutations in R20291, an epidemic RT012 isolate, VPI 10463 (ATCC 43255), a high toxin-producing RT087 strain, and 5325, representing the non-motile, agriculturally-associated 078 ribotype. To accomplish this, we used constructs that were previously successful in the 630 background to generate either a nonpolar *rstA* mutation via the CRISPR/cas9 system or an insertional disruption within the coding region of *rstA* (13, 18). An *rstA* mutant, in which the entire open reading frame was deleted, was generated in R20291 and confirmed by PCR (**Fig. S1, S2A**).

Despite the absence of reports that VPI 10463 accepts DNA (11), we were able to successfully conjugate exogenous DNA into VPI 10463 using a previously published heat shock method (43). However, these strains grew poorly and did not survive on plates longer than 48 h, resulting in few viable colonies throughout screening and/or selection and no detectable *rstA* mutants. Multiple 5325 *rstA* mutants were identified upon initial screenings, but this deletion was not stable after a single passage in any tested condition. Both VPI 10463 and 5325 were used throughout the rest of this study, as VPI 10463 is frequently used in laboratory settings and in the mouse model of *C. difficile* infection (44-48), and 5325 represents an emerging, clinically-relevant ribotype with unique characteristics (49, 50).

The amino acid sequence of RstA is identical in 630Δ*erm*, R20291, and VPI 10463, and only a single residue substitution is present in 5325 (V371I), suggesting that RstA function is highly conserved in these *C. difficile* strains. As RstA promotes the initiation of sporulation in 630Δ*erm* (18), we asked whether RstA exerts the same effect in strain R20291. Sporulation was measured after 24 h of growth on 70:30 sporulation agar by enumerating ethanol-resistant spores and viable vegetative cells, and by phase contrast microscopy. As expected, sporulation frequency was reduced ∼27-fold in the R20291 *rstA* mutant compared to the R20291 parent (**Fig. 1A**), indicating that RstA retains similar regulatory function in sporulation in 630Δ*erm* and R20291. The *rstA* mutation was complemented by integrating the *rstA* allele driven by its native promoter onto the chromosome via Tn*916* (51, 52; **Fig. S2B**), and the sporulation frequency in this strain was increased to greater than wild-type levels (**Fig. S2C**). VPI 10463 exhibited a relatively low sporulation rate of 6.3%, while we observed a robust sporulation frequency of 79.4% in 5325 (**Fig. 1A**).

**Figure 1.**
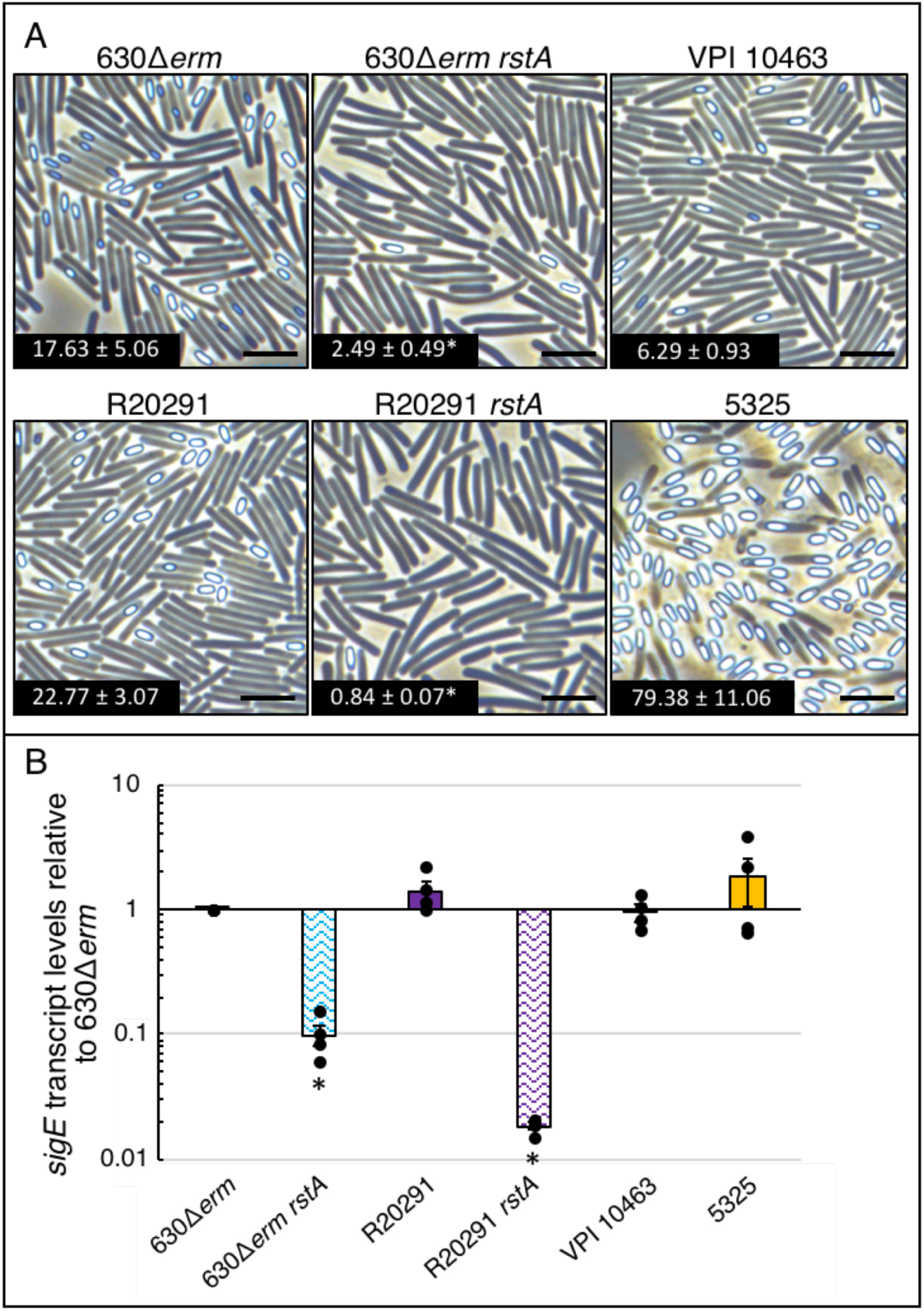
RstA promotes sporulation and Spo0A-dependent gene expression. (**A**) Representative phase contrast micrographs of 630Δ*erm*, 630Δ*erm rstA* (MC1118), R20291, R20291 *rstA* (MC1402), VPI 10463 and 5325 grown on 70:30 sporulation agar at H_24_. The numbers represent the percentage of ethanol-resistant spores to the total number of viable cells, at H_24_, and the standard error of the means of at least three biological replicates. The scale bar is equivalent to 5 *µ*m. (**B**) qRT-PCR analysis of *sigE* transcript levels in 630Δ*erm*, 630Δ*erm rstA* (MC1118), R20291, R20291 *rstA* (MC1402), VPI 10463 and 5325 grown on 70:30 sporulation agar at H_12_, relative to 630Δ*erm*. The means and standard error of the means for four biological replicates are shown, and markers represent the independent values within each mean. \**P* < 0.05 using a Student’s *t-*test comparing each *rstA* mutant to its isogenic parent.

To further probe the effect of RstA on sporulation initiation in *C. difficile*, we measured transcript levels of the Spo0A-dependent gene, *sigE*, which encodes an early-stage sporulation-specific sigma factor, using qRT-PCR (**Fig. 1B**). Relative expression of *sigE* in each strain mirrored the sporulation frequencies observed, indicating that RstA exerts its regulatory effect on the early stages of R20291 sporulation, as previously observed in 630Δ*erm* (18).

### RstA inhibits toxin gene expression and production in R20291

RstA directly represses *tcdA, tcdB, tcdR* and *sigD* transcription in 630Δ*erm* (13). The direct inhibition of two toxin gene activators, TcdR and SigD, as well as the direct repression of the toxin genes, results in a robust regulatory network that tightly controls toxin production (13). To determine whether RstA regulates toxin production in R20291, we first measured TcdA and TcdB protein levels present in TY culture supernatants after 24 h of growth via ELISA. As anticipated, toxin levels were increased approximately 11-fold in the 630Δ*erm rstA* mutant compared to its isogenic parent and were approximately 15-fold greater in the R20291 *rstA* mutant compared to its parent strain (**Fig. 2A**). Supernatant toxin levels in the R20291 *rstA* Tn*916*::*rstA* complemented strain returned to wild-type levels (**Fig. S2D**). Unsurprisingly, VPI 10463 exhibited high quantities of TcdA and TcdB in the culture supernatant, while the 5325 supernatant contained toxin levels comparable to 630Δ*erm* (**Fig. 2A**).

**Figure 2.**
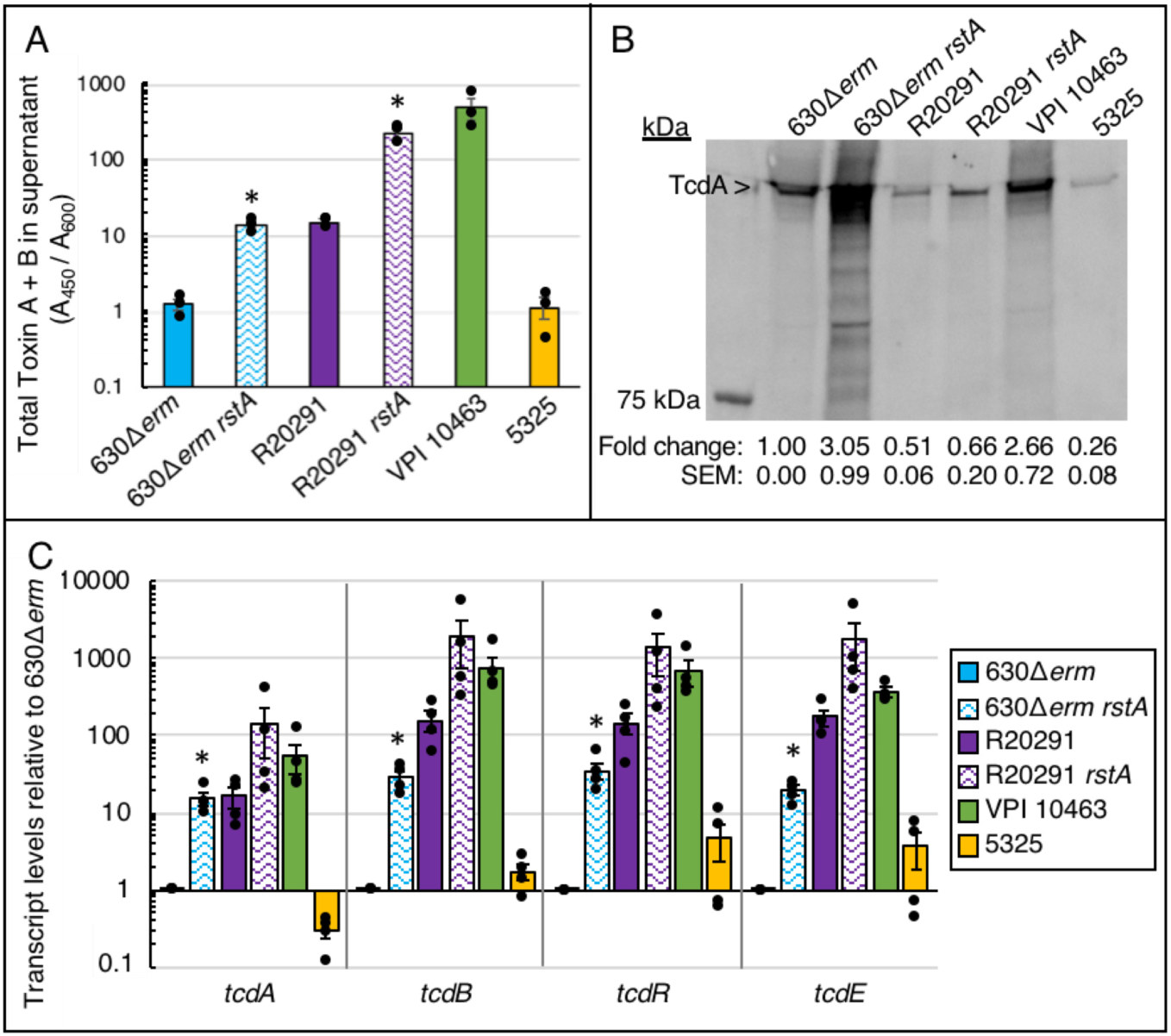
RstA inhibits *C. difficile* toxin production. (**A**) TcdA and TcdB present in the supernatants of 630Δ*erm*, 630Δ*erm rstA* (MC1118), R20291, R20291 *rstA* (MC1402), VPI 10463 and 5325 grown in TY medium, pH 7.4 at H_24_ were quantified by ELISA, as detailed in the Materials and Methods, and read as absorbance at 450 nm normalized to cell density (OD_600_). (**B**) Representative TcdA western blot analysis of 630Δ*erm*, 630Δ*erm rstA* (MC1118), R20291, R20291 *rstA* (MC1402), VPI 10463 and 5325 cells grown in TY medium, pH 7.4 at H_24_. (**C**) qRT-PCR analysis of *tcdA, tcdB, tcdR*, and *tcdE* transcript levels in 630Δ*erm*, 630Δ*erm rstA* (MC1118), R20291, R20291 *rstA* (MC1402), VPI 10463 and 5325 grown in TY medium, pH 7.4 at T_4._ (defined as four hours after entry into stationary phase). The means and standard error of the means for at least three biological replicates are shown, and markers represent the independent values within each mean. \**P* < 0.05 using a Student’s *t-*test comparing each *rstA* mutant to its isogenic parent.

To examine the amount of toxin within cells, western blot analysis were performed and TcdA protein levels were measured after 24 h of growth in TY medium. The western blot results indicated that the 630Δ*erm rstA* mutant had greater levels of TcdA relative to the parent strain, as previously observed (13, 18). R20291 cells contained approximately half the total amount of toxin than was found within 630Δ*erm* cells (**Fig. 2B**). The R20291 *rstA* mutant had a modest increase in TcdA toxin levels compared to R20291, and this effect was restored to wild-type level in the complemented strain (**Fig. S2E**). These data suggest that the majority of toxin synthesized by R20291 is secreted into the supernatant, whereas a lower ratio of the total 630Δ*erm* toxin is located extracellularly. There was ∼2.6-fold greater TcdA protein observed in VPI 10463 cells, while the 5325 cells had ∼4-fold less TcdA protein, compared to strain 630Δ*erm* (**Fig. 2B**).

To verify that RstA affected toxin production through regulation of toxin gene expression, we measured *tcdA, tcdB* and *tcdR* transcript levels from cultures grown in TY medium at T_4_, which corresponds to four hours after the cultures enter stationary phase, using qRT-PCR (**Fig. 2C**). As previously noted, *tcdA, tcdB* and *tcdR* transcript levels were increased ∼15 to 30-fold in the 630Δ*erm rstA* mutant compared to the 630Δ*erm* parent (13; **Fig. 2C**). Similar to the TcdA/TcdB ELISA results, the 630Δ*erm rstA* mutant and the R20291 strain had comparable levels of *tcdA* transcript; however, R20291 had greater transcript levels *tcdB* and *tcdR*. Transcript levels for all three *tcd* genes were increased ∼9-12-fold in the R20291 *rstA* mutant compared to its isogenic parent, although more variability was observed in the R20291 *rstA* mutant than in all other strains. VPI 10463 exhibited greater toxin gene expression than R20291, as previously observed in other conditions (22, 23). Of note, the R20291 *rstA* mutant had greater toxin gene expression than VPI 10463 (**Fig. 2C**), suggesting that R20291 is capable of producing high levels of toxin if RstA-dependent toxin inhibition is removed.

Because of the differences observed for toxin levels in the supernatant versus whole cell lysates, we asked whether RstA influenced gene expression of the toxin holin, *tcdE*, which is purported to play a role in toxin secretion in a strain-dependent manner (8, 29, 53). *tcdE* is the third gene encoded within the PaLoc (*tcdR-tcdB-tcdE-tcdA*), and its transcription is driven by its own, unmapped promoter (29). Transcript levels of *tcdE* showed a similar pattern of expression as the other toxin genes, with increased levels in both the 630Δ*erm rstA* and R20291 *rstA* strains compared to their isogenic parents (**Fig. 2C**). To determine whether the *tcdE* sequences in these isolates match the annotated Genbank sequences and encode a functional TcdE protein, primers flanking the *tcdE* open reading frame were used to amplify this region, and the resulting PCR products were purified and sequenced (data not shown). The 630Δ*erm*, R20291 and VPI 10463 *tcdE* sequences were identical to the annotated sequences. These *tcdE* sequences differ only in a nonsynonymous mutation in the R20291 sequence, resulting in a conserved I151V amino acid substitution. Analysis of M120, a sequenced RT078 reference genome, revealed the absence of the M1 start codon, leaving only the M25 and M27 start codons present. The *tcdE* open reading frame is capable of producing three isoforms, with TcdE_142_, initiating from M25, as the predominant isoform produced (29). TcdE_142_ also exhibits the greatest holin activity (29). TcdE_166_, the full-length isoform initiated from M1, is not efficiently translated *in vivo*, and the degeneration of the M1 start codon in the RT078 strains supports the hypothesis that this isoform is not essential for *C. difficile* toxin secretion. Altogether, these data suggest that even though RstA inhibits *tcdE* gene expression, RstA does not contribute to the strain-dependent differences observed for TcdE-mediated toxin secretion.

In strain 630Δ*erm*, RstA inhibits *tcdA* and *tcdB* transcription directly by binding their respective promoters (13). To determine whether RstA inhibits *tcdA* and *tcdB* transcription in R20291, alkaline phosphatase reporters were constructed by fusing the promoter regions of these genes to *phoZ* and integrating the reporters into the chromosomes of R20291 and R20291 *rstA*. P*tcdA*::*phoZ* and P*tcdB*::*phoZ* reporter activity was increased approximately 13- and 10-fold, respectively, in the R20291 *rstA* background (**Fig. 3**), indicating that RstA represses *tcdA* and *tcdB* transcription. The σ^TcdR^-dependent promoters of *tcdA* and *tcdB* and the putative RstA binding sites, which overlap the -35 consensus sequences of these promoters, are identical in 630Δ*erm* and R20291, strongly suggesting that RstA directly represses toxin gene transcription similarly in both strains. Additionally, as observed in the qRT-PCR analysis of these same genes (**Fig. 2C**), reporter activity from the toxin gene promoter fusions in the R20291 *rstA* background was highly variable (**Fig. 3**), suggesting that the absence of RstA enhances the variability of *tcdR, tcdA*, and *tcdB* transcription (see below).

**Figure 3.**
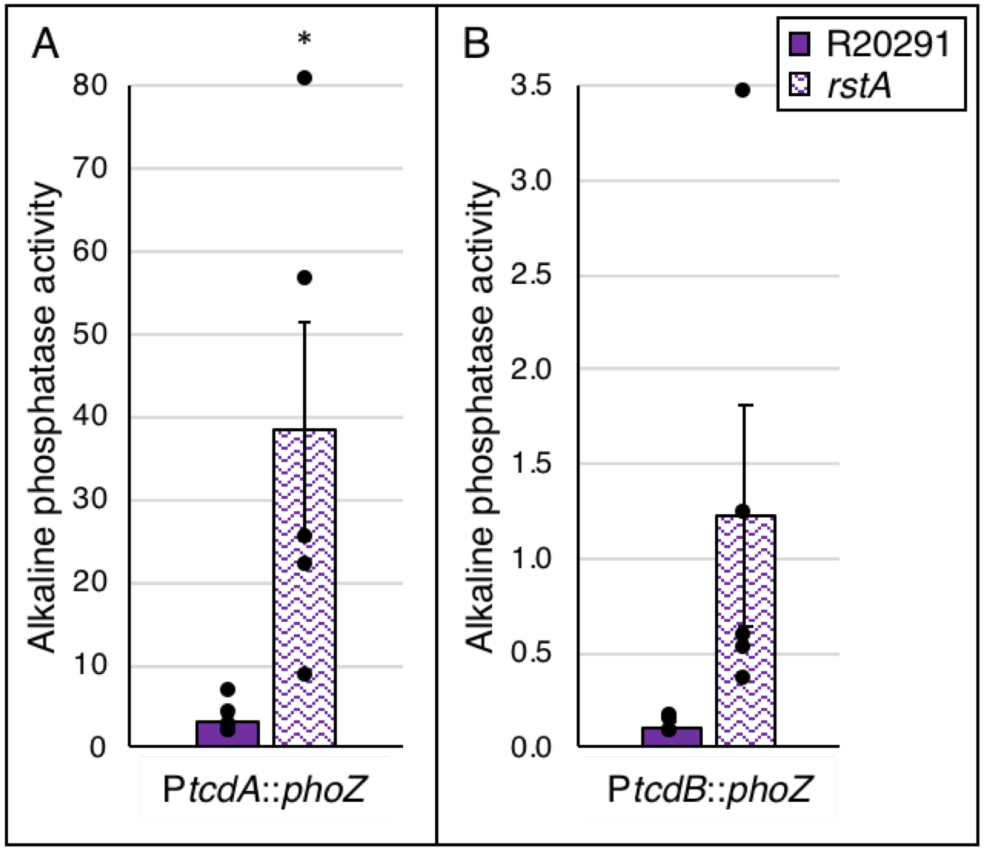
RstA inhibits *tcdA* and *tcdB* transcription in R20291. Alkaline phosphatase (AP) activity of (**A**) P*tcdA*::*phoZ* and (**B**) P*tcdB*::*phoZ* integrated into the chromosome using Tn*916* in R20291 (MC1451 and MC1453) or R20291 *rstA* (MC1452 and MC1454). The means and standard error of the means are shown for five biological replicates. \**P* < 0.05 using a Student’s *t-*test.

Expression of *tcdR* is driven by several promoters (**Fig. 4A**), one of which requires the motility-specific sigma factor, SigD (54, 55). RstA inhibits *sigD* gene expression, the fourth gene encoded within the *flgB* operon, by directly binding to the *flgB* promoter (13). Interestingly, neither *flgB* nor *sigD* transcript levels are significantly altered in the R20291 *rstA* mutant compared to the R20291 parent (**Fig. S3**). SigD activity is slightly greater in the R20291 *rstA* mutant, as evidenced by increased transcript levels of the SigD-dependent gene, *fliC*, compared to the R20291 parent strain (**Fig. S3**), although this regulatory effect is not statistically significant. Strain 5325 was omitted from this analysis, as its genome is lacking the flagellar genes. Previously, reporter fusions revealed that RstA represses PflgB^(630Δ*erm*)^ and P*flgB*^(R20291)^ activity ∼1.5-fold between the 630Δ*erm* and 630Δ*erm rstA* strains (13). Further, biotin-pulldown assays with 630Δ*erm* lysates showed that RstA directly binds to the R20291 *flgB* promoter (13). These previous studies were all performed using the 630Δ*erm* background. Our current data demonstrates no significant effect by RstA on motility gene transcription in the R20291 background, suggesting that there are significant strain-specific differences in RstA-dependent regulation of *flgB* transcription between the 630Δ*erm* and R20291 backgrounds. RstA may control toxin production in R20291 by influencing SigD activity, but there do not appear to be strong, direct effects of RstA on *sigD* gene expression in this strain.

**Figure 4.**
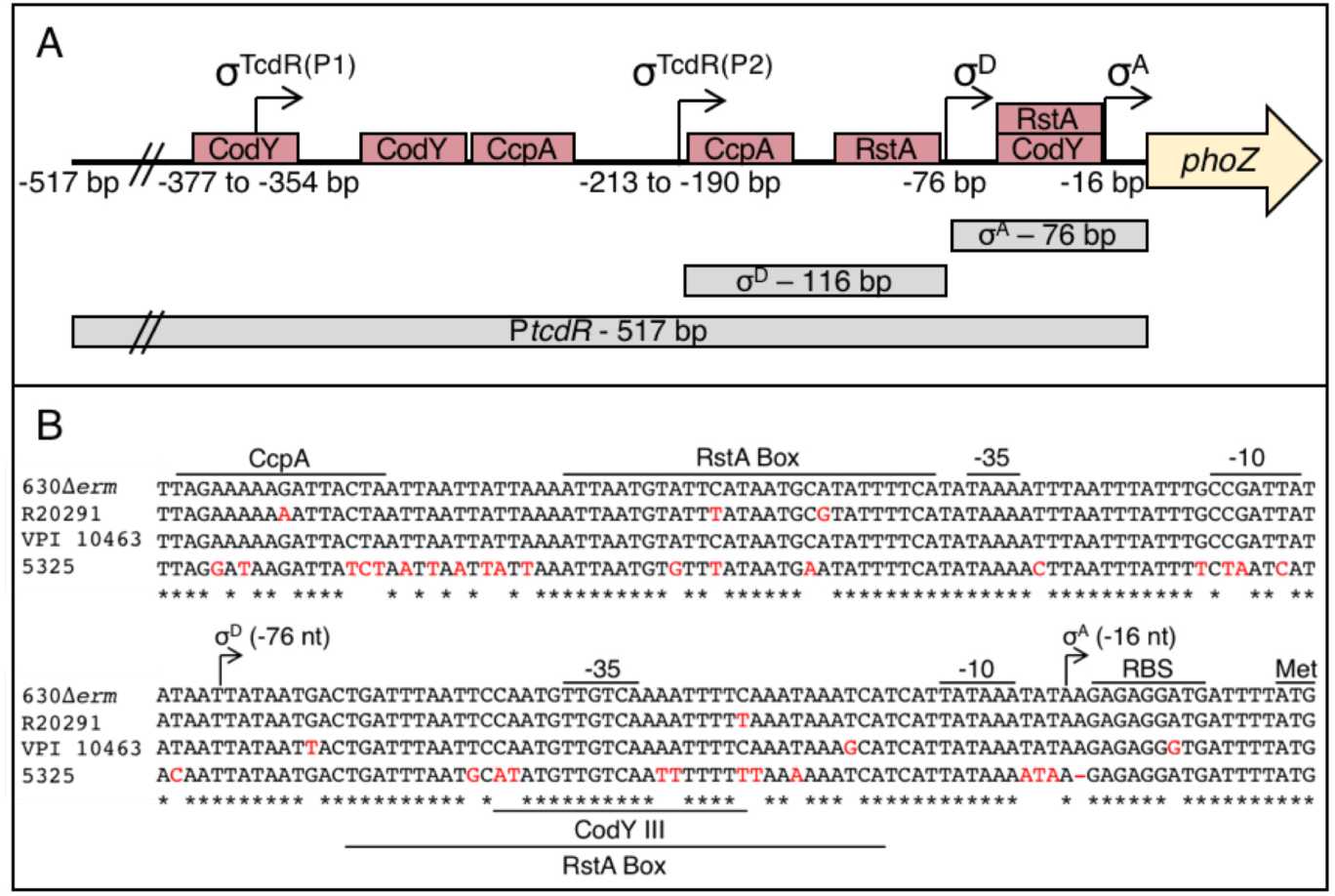
Strain-specific nucleotide changes overlap key regulatory recognition sites within the *tcdR* promoter region. (**A**) Schematic depicting the mapped transcriptional start sites (bent arrows; the nucleotide position is marked below, and the specific sigma-factor that recognizes the promoter is denoted above) and the RstA, CodY and CcpA binding sites (red boxes) of 630Δ*erm* (11-13). The sequence composition of the three reporter fusions created for each strain are indicated as gray boxes below. (**B**) Alignment and annotation of nucleotide changes within the σ^D^ - and σ^A^ - dependent *tcdR* promoter regions for four diverse *C. difficile* strains including 630Δ*erm*, R20291, VPI 10463, and 5325. Specific sequence features are identified, and non-conserved nucleotides are marked in red.

### RstA and CodY regulate *tcdR* transcription in concert

Our previous work revealed that RstA controls 630Δ*erm tcdR* transcription by directly binding to the σ^A^- and σ^D^-dependent promoters directly upstream of *tcdR* (13; **Fig. 4A**). P*tcdR*(σ^D^) features an imperfect inverted repeat immediately upstream of the conserved -35 sequence that likely serves as the RstA-binding site (13; **Fig. 4A**). Notably, the RstA box and CodY III box of P*tcdR*(σ^A^) perfectly overlap each other, suggesting that inhibiting P*tcdR*(σ^A^) transcription is important for toxin regulation. Alignment of the P*tcdR* regions from 630Δ*erm*, R20291, VPI 10463 and 5325 revealed multiple single nucleotide substitutions throughout the conserved regulatory features (**Fig. 4B**). The R20291 and VPI 10463 P*tcdR*(σ^A^) sequences each revealed distinct single nucleotide changes within the RstA box and CodY III binding site of P*tcdR*(σ^A^). Additionally, the 5325 sequence contains eight individual nucleotide changes within this region (five mismatches within the CodY III binding site and an additional three extending through the RstA Box). As 5325 is non-motile and the flagellar genes, including *sigD*, are absent from its genome, we accordingly noted significant sequence degeneration in the region that aligns with 630Δ*erm* P*tcdR*(σ^D^) and hypothesized that the 5325 P*tcdR* region does not rely on SigD for toxin gene regulation. Further, we hypothesized that the nucleotide changes in each strain alter RstA and CodY regulation of *tcdR* expression. To test these hypotheses, the full-length *tcdR* promoter and the fragments corresponding to the σ^A^- and σ^D^-dependent promoters from R20291, VPI 10463 and 5325 were amplified and cloned upstream of the *phoZ* gene in a multi-copy plasmid. We did not include an individual P*tcdR*(σ^D^)^(VPI 10463)^ reporter construct, as the 630Δ*erm* and VPI 10463 sequences spanning this region are identical (**Fig. 4B**). All of the reporter constructs were expressed in the 630Δ*erm*, 630Δ*erm rstA* and 630Δ*erm codY* backgrounds to eliminate additional strain-dependent regulatory effects. Alkaline phosphatase reporter assays were performed on cells containing promoter::*phoZ* fusions from samples collected after 24 h growth in TY medium.

The reporter activity for the full-length *tcdR* promoters expressed in the 630Δ*erm* background were equivalent, except the full-length P*tcdR*^(R20291)^ fusion exhibited ∼1.7-fold lower activity compared to the others (**Fig. 5A**). Interestingly, the activities of the R20291, VPI 10463 and 5325 P*tcdR*(σ^A^) reporters in the 630Δ*erm* background were ∼2.4- to 3.6-fold higher than the activity from the P*tcdR*(σ^A^)^(630Δ*erm*)^ fusion (**Fig. 5A**), suggesting that CodY and RstA do not repress basal transcription from these promoters as efficiently. These data indicate that the CodY III and RstA consensus sequences in the 630Δ*erm* promoter allow for the greatest repression by RstA and CodY, and that nucleotide changes within this region reduce the repressive effect by RstA and/or CodY.

**Figure 5.**
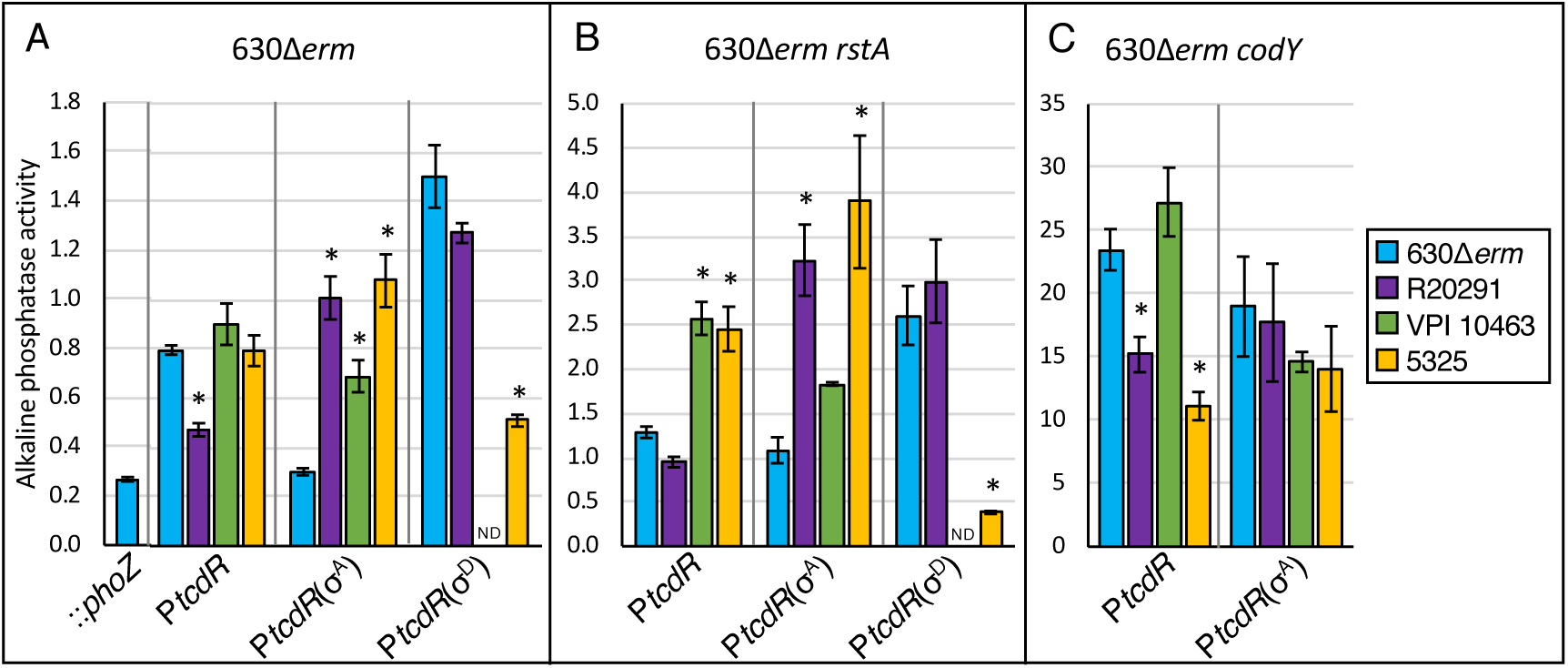
Strain-specific nucleotide changes within the *tcdR* promoter affect RstA- and CodY-dependent regulation. Alkaline phosphatase (AP) activity of a series of P*tcdR*::*phoZ* fusions, expressed from a plasmid in 630Δ*erm* (**A**), 630Δ*erm rstA* (**B**), and 630Δ*erm codY*::*erm* (**C**) comprised of the full-length *tcdR* promoter from 630Δ*erm* (MC1088/MC1330/MC1552), R20291 (MC1458/MC1459/MC1554), VPI 10463 (MC1563/MC1564/MC1558), and 5325 (MC1425/MC1426/MC1556), the σ^A^ –dependent *tcdR* promoter from 630Δ*erm* (MC1285/MC1331/MC1553), R20291 (MC1460/MC1461/MC1555), VPI 10463 (MC1565/MC1566/MC1559), and 5325 (MC1427/MC1428/MC1557), and the σ^D^-dependent *tcdR* promoter from 630Δ*erm* (MC1145/MC1332), R20291 (MC1462/MC1463), and 5325 (MC1429/MC1430). As the nucleotide sequences of the 630Δ*erm* and VPI 10463 *tcdR*(σ^D.^) promoters are identical, an individual P*tcdR*(σ^D^) fusion for VPI 10463 was not constructed (ND). The promoterless ::*phoZ* reporter carried by 630Δ*erm* (MC448) was included as a negative control. Strains were grown in TY medium, pH 7.4, supplemented with 2 *µ*g ml^−1^ thiamphenicol, and samples assayed for alkaline phosphatase activity were collected at H_24_. The means and standard error of the means of at least three biological replicates are shown. \**P* < 0.05 using one-way ANOVA followed by Dunnett’s multiple comparisons test compared to the PtcdR^(630Δerm)^ -derived promoter within the same set. The activity of each reporter in the 630Δ*erm rstA* background or the 630Δ*erm codY* background compared to 630Δ*erm*, with the exception of the P*tcdR*(σ^D^)^(5325)^ reporter, was statistically significant (*P* < 0.05) using one-way ANOVA followed by Dunnett’s multiple comparisons test (not shown in figure).

The reporter activities from the 630Δ*erm*-derived promoter fusions expressed in the *rstA* deletion mutant were similar to those previously reported (13), with the strongest RstA-dependent effects (∼3.5 fold) observed for the P*tcdR*(σ^A^) reporter, and ∼1.6- to 1.7-fold changes in reporter activity for the full-length and the P*tcdR*(σ^D^) reporters (**Table 3; Fig. 5A** and **B**). The full-length *tcdR* and P*tcdR*(σ^A^) reporters from R20291, VPI 10463 and 5325 and the P*tcdR*(σ^D^)^(R20291)^ reporter showed ∼2-3-fold higher activity when expressed in the 630Δ*erm rstA* mutant compared to the parent strain (**Table 3**; **Fig. 5B**). These data demonstrate RstA-dependent repression of all strains’ *tcdR* promoters. Minor changes in activity were observed for the four different *tcdR* promoters, but none of the sequence changes fully relieved RstA-mediated repression, suggesting that RstA inhibition of *tcdR* transcription is conserved in *C. difficile.* As expected, the fusion containing the unconserved P*tcdR*(σ^D^)^(5325)^exhibited no reporter activity above background levels, consistent with the degenerative sequence.

**Table 3.** Fold changes in P*tcdR* reporter activity expressed in the 630Δ*erm*, 630Δ*erm rstA* or 630Δ*erm codY* backgrounds.

| Reporter Fusion | Fold change in reporter activity |  |
| --- | --- | --- |
| | *rstA* / 630Δ*erm*<sup>a</sup> | *codY* / 630Δ*erm*<sup>a</sup> |
| *PtcdR*<sup>(630Δ*erm*)</sup>::*phoZ* | 1.61 | 29.5 |
| *PtcdR*<sup>(R20291)</sup>::*phoZ* | 2.01 | 32.2 |
| *PtcdR*<sup>(VPI 10463)</sup>::*phoZ* | 2.85 | 30.2 |
| *PtcdR*<sup>(5325)</sup>::*phoZ* | 3.09 | 14.0 |
| *PtcdR*($\sigma^A$)<sup>(630Δ*erm*)</sup>::*phoZ* | 3.59 | 63.0 |
| *PtcdR*($\sigma^A$)<sup>(R20291)</sup>::*phoZ* | 3.20 | 17.6 |
| *PtcdR*($\sigma^A$)<sup>(VPI 10463)</sup>::*phoZ* | 2.65 | 21.2 |
| *PtcdR*($\sigma^A$)<sup>(5325)</sup>::*phoZ* | 3.61 | 13.0 |
| *PtcdR*($\sigma^D$)<sup>(630Δ*erm*)</sup>::*phoZ* | 1.74 | ND<sup>b</sup> |
| *PtcdR*($\sigma^D$)<sup>(R20291)</sup>::*phoZ* | 2.35 | ND<sup>b</sup> |
| *PtcdR*($\sigma^D$)<sup>(5325)</sup>::*phoZ* | 0.75 | ND<sup>b</sup> |

<sup>a</sup>Fold change of the average reporter activity (presented in Fig. 5) in the 630Δ *erm* *rstA* or 630Δ *erm* *codY* strain divided by the activity in the 630Δ *erm* parent strain.
<sup>b</sup>ND, not determined; *PtcdR*( $\sigma^D$ ) reporter activity was not tested in the 630Δ *erm* *codY* as no CodY binding site is located within the *tcdR* $\sigma^D$ -dependent promoter region (11).

To assess the specific repression CodY exerts on *tcdR* transcription, we compared the activity of the full-length *tcdR* and the σ^A^-dependent promoter reporters in 630Δ*erm* and 630Δ*erm codY*. Promoter activity from the 630Δ*erm*, R20291 and VPI 10463 full-length P*tcdR* reporters were similarly elevated in the *codY* mutant, relative to the parent strain (∼30-fold; **Table 3**). However, the P*tcdR*^(5325)^ reporter exhibited approximately half the activity in the absence of CodY (14-fold change; **Table 3**; **Fig. 5C**). Our data suggest that CodY derepression of *tcdR* expression is weaker in 5325, which corresponds to the greater number of mismatches within the CodY III box of the 5325 sequence (**Fig. 4B**). We also observed that activity of the 630Δ*erm*, R20291, VPI 10463 and 5325 P*tcdR*(σ^A^) reporters were similar in the 630Δ*erm codY* mutant (**Fig. 5C**). But, the lower activity from the P*tcdR*(σ^A^)^(630Δ*erm*)^ reporter in the 630Δ*erm* background resulted in a 63-fold change in activity when measured in the 630Δ*erm codY* mutant (**Table 3**), emphasizing that CodY most efficiently represses transcription from the 630Δ*erm*-derived σ^A^-dependent promoter, as it contains the most conserved CodY binding site. Overall, relief of CodY repression results in greater activity from the P*tcdR* reporters than the absence of RstA (**Fig. 5B** versus **5C**). However, as these regulators respond to different cofactors, RstA and CodY repression likely prevents full *tcdR* transcription unless DNA-binding by both regulators is relieved.

### RstA autoregulates its transcription in R20291

RstA directly represses its own transcription in 630Δ*erm* by binding to an imperfect inverted repeat that overlaps the -10 consensus sequence and start of transcription for the σ^A^-dependent *rstA* promoter (13, 18; **Fig. S4**). This direct negative autoregulation results in constitutive expression of *rstA* throughout growth and is typical of the RRNPP family of proteins (18, 56, 57). Because the *rstA* mutations we generated were clean deletions and eliminated the ability to detect *rstA* transcripts (**Fig. S5A** and **B**), we constructed reporter fusions comprising the 500 bp upstream from the *rstA* translational start from both 630Δ*erm* and R20291, to test whether RstA represses *rstA* transcription in the R20291 background (**Fig. S4**; herein referred to as P*rstA*^(630Δ*erm*)^::*phoZ* and P*rstA*^(R20291)^::*phoZ*). This upstream region includes the 5’ end of the open reading frame of the divergently transcribed upstream gene, *CD3669*, and this region contains three nucleotide substitutions that are located in the R20291 sequence compared to 630Δ*erm* (**Fig. S4**). To determine how RstA impacts its own transcription in these two strains, the fusions were integrated into their respective parent and *rstA* mutant chromosomes, and the P*rstA*^(R20291)^::*phoZ* reporter was also integrated into the 630Δ*erm* and 630Δ*erm rstA* mutant strains to allow for direct comparisons.

After eight hours of growth on 70:30 sporulation agar, we found that promoter activity from the P*rstA*^(630Δ*erm*)^::*phoZ* reporter was ∼1.3-fold greater in the 630Δ*erm rstA* background compared to the parent strain (**Fig. 6**), which correlates with the fold changes seen from the same fusion on a plasmid expressed in 630Δ*erm* and an isogenic *rstA*::*erm* mutant (13, 18). Surprisingly, P*rstA*^(R20291)^::*phoZ* exhibited identical activity in both the 630Δ*erm* and 630Δ*erm rstA* backgrounds (**Fig. 6**), indicating that there are modest sequence-dependent effects influencing expression from the 630Δ*erm* and R20291 *rstA* promoters. Finally, the P*rstA*^(R20291)^::*phoZ* reporter exhibited a significant reduction in activity in the R20291 parent background compared to the 630Δ*erm* parent background and a ∼2.7-fold increase in activity in the *rstA* mutant compared to the R20291 parent (**Fig. 6**). We observed significantly higher *rstA* transcript levels in the R20291 strain compared to 630Δ*erm* in two different growth media (**Fig. S5A** and **B**), suggesting that basal expression of *rstA* is higher in R20291 and may explain some of the strain-dependent differences observed with RstA regulation. Altogether, these data indicate that RstA strongly represses its own expression in the R20291 background and reveal the presence of strain-dependent differences that influence RstA autoregulation.

**Figure 6.**
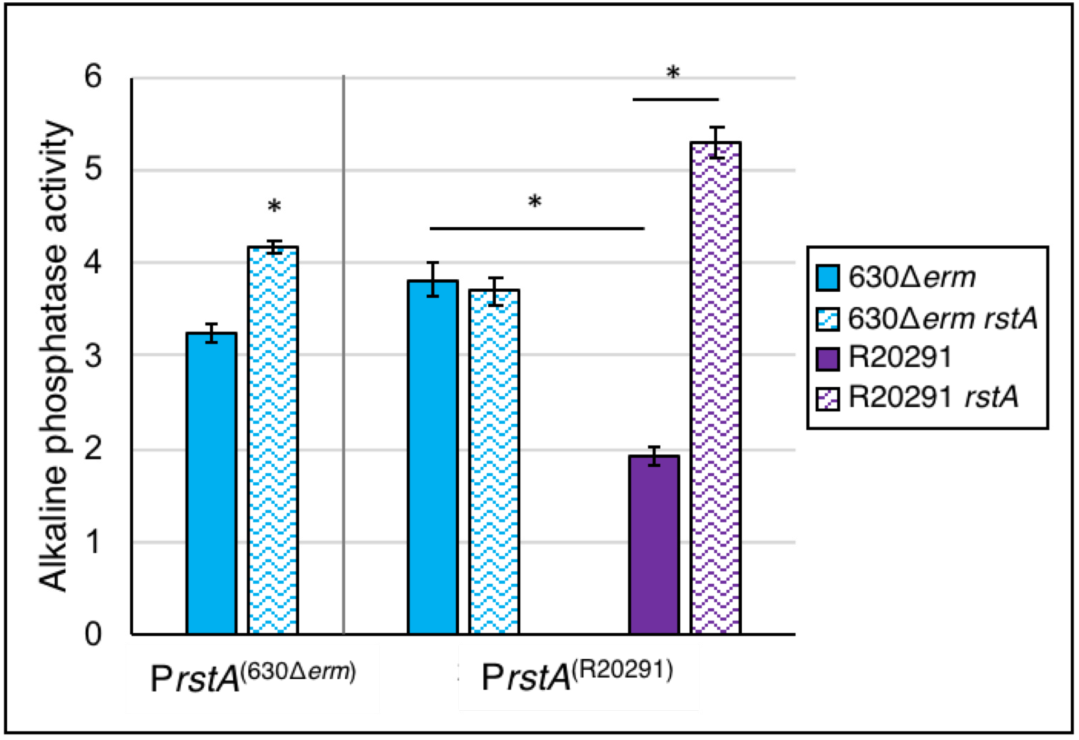
RstA negatively autoregulates its own expression in R20291. Alkaline phosphatase (AP) activity of the *rstA* promoter from 630Δ*erm* (P*rstA*^(630Δ*erm*)^ ::*phoZ*) integrated into 630Δ*erm* (MC1640) and 630Δ*erm rstA* (MC1641), or the *rstA* promoter from R20291 (P*rstA*^(R20291)^ ::*phoZ*) integrated into 630Δ*erm* (MC1668), 630Δ*erm rstA* (MC1669), R20291 (MC1642) and R20291 *rstA* (MC1643). Strains were grown on 70:30 sporulation agar, and samples assayed for alkaline phosphatase activity were collected at H_8_. The means and standard error of the means of four biological replicates are shown. \**P* < 0.05 using a Student’s *t-*test for the P*rstA*^(630Δ*erm*)^::*phoZ* comparison or a one-way ANOVA followed by the Tukey’s multiple comparisons test for the P*rstA*^(R20291)^::*phoZ* comparisons.

## DISCUSSION

*C. difficile* toxin production is the mediator of significant gastrointestinal disease throughout the course of infection in the host (1). The production of toxin has been attributed to nutritional availability, temperature control, activity of the flagellar-specific sigma factor, SigD, and the synthesis of secondary messengers (54, 55, 58-62, reviewed in 63), yet strain-specific differences have been extensively noted for most aspects of toxin gene expression, synthesis and secretion (9, 10, 22, 23, 25, 29, 64). Further, many of these environmental conditions also affect another important aspect of *C. difficile* pathogenesis: sporulation. The multifunctional regulator, RstA, supports early stage sporulation events and directly inhibits toxin gene expression in response to a putative cofactor in the historical strain, 630Δ*erm* (13, 18). In this study, we determined the regulatory effects RstA has on toxin production and sporulation in the epidemic isolate, R20291, and assessed the effects of RstA and CodY inhibition of the *tcdR* promoters from four diverse *C. difficile* strains.

As in the 630 background, we found that RstA induces R20291 sporulation and inhibits *tcdR, tcdA* and *tcdB* expression. Surprisingly, motility gene expression was not strongly affected, suggesting that the effect of RstA on SigD-dependent toxin gene expression is less pronounced in the R20291 background. Unfortunately, attempts to assess the impact of RstA on the individual R20291 P*tcdR* promoters were unsuccessful due to the undetectable level of alkaline phosphatase activity from the single-copy reporter in the R20291 and R20291 *rstA* backgrounds; however, the remarkably low expression of the *tcdR* promoter is well documented (13, 55, 59, 65, 66). Further studies to elucidate the regulatory effects of RstA-dependent toxin gene repression in R20291 and additional *C. difficile* strains will illuminate the seemingly minor differences in the toxin regulatory network between strains that may significantly impact total toxin production and the resulting virulence.

One apparent difference in the regulation of toxin by RstA between the R20291 and 630Δ*erm* strains is the inherent variability in toxin gene expression (**Figs. 2C** and **3**). The increased variability of toxin gene expression in the R20291 *rstA* mutant may be a result of bistable toxin production mediated by TcdR, in which individual cells differentially express toxin genes as either toxin-ON or toxin-OFF (66). Although our data are representative of the population, the higher incidence of variability in *tcdA, tcdB, tcdR* and *tcdE* transcript levels and in *tcdA*::*phoZ* and *tcdB*::*phoZ* reporter activity in the R20291 *rstA* mutant implicate RstA as a mediator of bimodal toxin gene expression by biasing the population in the toxin-OFF state, as is the case for CodY (66). This variability in toxin gene expression is not observed in the 630Δ*erm rstA* mutant, corroborating previous results showing that a higher proportion of cells are toxin-ON in 630Δ*erm*, likely due to higher *sigD* expression (66, 67).

The regulatory effects by RstA appeared more marked in the R20291 strain compared to the previously studied 630Δ*erm* background, with a more severe sporulation defect than in 630Δ*erm* (7-fold in 630Δ*erm* versus 27-fold in R20291; **Fig. 1**). In addition, greater fold-changes in activity were observed with the P*rstA*, P*tcdA* and P*tcdB* reporters in the R20291 and R20291 *rstA* strains, than observed in the corresponding 630Δ*erm* constructs (**Figs 3 and 6**). The *rstA* transcript levels were also greater in R20291 compared to 630Δ*erm* (**Fig. S5**), suggesting that higher levels of RstA influence RstA activity and its regulatory effects. Further, our observations that VPI 10463 is a poor spore former, yet high toxin-producer and that 5325 produces prolific spores and low levels of toxin, suggests that RstA regulation may be involved in these opposing phenotypes.

The dual repressive effects of RstA and CodY on P*tcdR*(σ^A^)-dependent transcription highlight the importance of tightly regulating toxin production in *C. difficile*. Nucleotide substitutions within the RstA and CodY consensus sequences resulted in minor differences with P*tcdR* reporter activity (**Figs. 4** and **5**; **Table 3**), but none of the changes completely abolished repression by these regulators. Our data suggest that RstA and CodY-mediated repression of toxin gene expression occur in the R20291, VPI 10463 and 5325 strains, although caution in extrapolating this data is warranted, as the differences observed in P*rstA*^(R20291)^::*phoZ* activity between its native R20291 background and the 630Δ*erm* heterologous background were striking. These findings further underscore the importance of differentiating between sequence-specific and strain-specific regulatory impacts. Identifying the cofactor that regulates RstA DNA-binding activity and deciphering the molecular mechanisms by which RstA promotes sporulation will aid in dissecting these sequence-dependent and strain-dependent regulatory differences.

## ACKNOWLEDGEMENTS

We give special thanks to the members of McBride lab for helpful suggestions and discussions during the course of this work. This research was supported by the U.S. National Institutes of Health through research grants AI116933 and AI121684 to S.M.M. The content of this manuscript is solely the responsibility of the authors and does not necessarily reflect the official views of the National Institutes of Health.

## REFERENCES

1. Kuehne SA, Cartman ST, Heap JT, Kelly ML, Cockayne A, Minton NP. 2010. The role of toxin A and toxin B in Clostridium difficile infection. Nature 467:711–3.

2. Just I, Selzer J, Wilm M, von Eichel-Streiber C, Mann M, Aktories K. 1995. Glucosylation of Rho proteins by Clostridium difficile toxin B. Nature 375:500–3.

3. Just I, Wilm M, Selzer J, Rex G, von Eichel-Streiber C, Mann M, Aktories K. 1995. The enterotoxin from Clostridium difficile (ToxA) monoglucosylates the Rho proteins. Journal of Biological Chemistry 270:13932–13936.

4. Chumbler NM, Farrow MA, Lapierre LA, Franklin JL, Lacy DB. 2016. Clostridium difficile Toxins TcdA and TcdB Cause Colonic Tissue Damage by Distinct Mechanisms. Infect Immun 84:2871–7.

5. Hammond GA, Johnson JL. 1995. The toxigenic element of Clostridium difficile strain VPI 10463. Microb Pathog 19:203–13.

6. Mani N, Dupuy B. 2001. Regulation of toxin synthesis in Clostridium difficile by an alternative RNA polymerase sigma factor. Proc Natl Acad Sci U S A 98:5844–9.

7. Tan KS, Wee BY, Song KP. 2001. Evidence for holin function of tcdE gene in the pathogenicity of Clostridium difficile. J Med Microbiol 50:613–9.

8. Govind R, Dupuy B. 2012. Secretion of Clostridium difficile toxins A and B requires the holin-like protein TcdE. PLoS Pathog 8:e1002727.

9. Matamouros S, England P, Dupuy B. 2007. Clostridium difficile toxin expression is inhibited by the novel regulator TcdC. Mol Microbiol 64:1274–88.

10. Cartman ST, Kelly ML, Heeg D, Heap JT, Minton NP. 2012. Precise manipulation of the Clostridium difficile chromosome reveals a lack of association between the tcdC genotype and toxin production. Appl Environ Microbiol 78:4683–90.

11. Dineen SS, Villapakkam AC, Nordman JT, Sonenshein AL. 2007. Repression of Clostridium difficile toxin gene expression by CodY. Mol Microbiol 66:206–19.

12. Antunes A, Martin-Verstraete I, Dupuy B. 2011. CcpA-mediated repression of Clostridium difficile toxin gene expression. Mol Microbiol 79:882–99.

13. Edwards AN, Anjuwon-Foster BR, McBride SM. 2019. RstA Is a Major Regulator of Clostridioides difficile Toxin Production and Motility. MBio 10.

14. Deakin LJ, Clare S, Fagan RP, Dawson LF, Pickard DJ, West MR, Wren BW, Fairweather NF, Dougan G, Lawley TD. 2012. The Clostridium difficile spo0A gene is a persistence and transmission factor. Infect Immun 80:2704–11.

15. Edwards AN, McBride SM. 2014. Initiation of sporulation in Clostridium difficile: a twist on the classic model. FEMS Microbiol Lett 358:110–8.

16. Nawrocki KL, Edwards AN, Daou N, Bouillaut L, McBride SM. 2016. CodY-Dependent Regulation of Sporulation in Clostridium difficile. J Bacteriol 198:2113–30.

17. Antunes A, Camiade E, Monot M, Courtois E, Barbut F, Sernova NV, Rodionov DA, Martin-Verstraete I, Dupuy B. 2012. Global transcriptional control by glucose and carbon regulator CcpA in Clostridium difficile. Nucleic Acids Res 40:10701–18.

18. Edwards AN, Tamayo R, McBride SM. 2016. A novel regulator controls Clostridium difficile sporulation, motility and toxin production. Mol Microbiol 100:954–71.

19. Dineen SS, McBride SM, Sonenshein AL. 2010. Integration of metabolism and virulence by Clostridium difficile CodY. J Bacteriol 192:5350–62.

20. Warny M, Pepin J, Fang A, Killgore G, Thompson A, Brazier J, Frost E, McDonald LC. 2005. Toxin production by an emerging strain of Clostridium difficile associated with outbreaks of severe disease in North America and Europe. Lancet 366:1079–84.

21. Akerlund T, Persson I, Unemo M, Noren T, Svenungsson B, Wullt M, Burman LG. 2008. Increased sporulation rate of epidemic Clostridium difficile Type 027/NAP1. J Clin Microbiol 46:1530–3.

22. Merrigan M, Venugopal A, Mallozzi M, Roxas B, Viswanathan VK, Johnson S, Gerding DN, Vedantam G. 2010. Human hypervirulent Clostridium difficile strains exhibit increased sporulation as well as robust toxin production. J Bacteriol 192:4904–11.

23. Vohra P, Poxton IR. 2011. Comparison of toxin and spore production in clinically relevant strains of Clostridium difficile. Microbiology 157:1343–53.

24. Burns DA, Heeg D, Cartman ST, Minton NP. 2011. Reconsidering the sporulation characteristics of hypervirulent Clostridium difficile BI/NAP1/027. PLoS One 6:e24894.

25. Mackin KE, Carter GP, Howarth P, Rood JI, Lyras D. 2013. Spo0A differentially regulates toxin production in evolutionarily diverse strains of Clostridium difficile. PLoS One 8:e79666.

26. Rupnik M, Janezic S. 2016. An Update on Clostridium difficile Toxinotyping. J Clin Microbiol 54:13–8.

27. Spigaglia P, Mastrantonio P. 2002. Molecular analysis of the pathogenicity locus and polymorphism in the putative negative regulator of toxin production (TcdC) among Clostridium difficile clinical isolates. J Clin Microbiol 40:3470–5.

28. Curry SR, Marsh JW, Muto CA, O’Leary MM, Pasculle AW, Harrison LH. 2007. tcdC genotypes associated with severe TcdC truncation in an epidemic clone and other strains of Clostridium difficile. J Clin Microbiol 45:215–21.

29. Govind R, Fitzwater L, Nichols R. 2015. Observations on the Role of TcdE Isoforms in Clostridium difficile Toxin Secretion. J Bacteriol 197:2600–9.

30. Stabler RA, He M, Dawson L, Martin M, Valiente E, Corton C, Lawley TD, Sebaihia M, Quail MA, Rose G, Gerding DN, Gibert M, Popoff MR, Parkhill J, Dougan G, Wren BW. 2009. Comparative genome and phenotypic analysis of Clostridium difficile 027 strains provides insight into the evolution of a hypervirulent bacterium. Genome Biol 10:R102.

31. Smith CJ, Markowitz SM, Macrina FL. 1981. Transferable tetracycline resistance in Clostridium difficile. Antimicrob Agents Chemother 19:997–1003.

32. Sorg JA, Dineen SS. 2009. Laboratory maintenance of Clostridium difficile. Curr Protoc Microbiol Chapter 9:Unit9A 1.

33. Edwards AN, Suarez JM, McBride SM. 2013. Culturing and maintaining Clostridium difficile in an anaerobic environment. J Vis Exp doi:10.3791/50787:e50787.

34. Putnam EE, Nock AM, Lawley TD, Shen A. 2013. SpoIVA and SipL are Clostridium difficile spore morphogenetic proteins. J Bacteriol 195:1214–25.

35. Luria SE, Burrous JW. 1957. Hybridization between Escherichia coli and Shigella. J Bacteriol 74:461–476.

36. Purcell EB, McKee RW, McBride SM, Waters CM, Tamayo R. 2012. Cyclic diguanylate inversely regulates motility and aggregation in Clostridium difficile. J Bacteriol 194:3307–16.

37. Childress KO, Edwards AN, Nawrocki KL, Woods EC, Anderson SE, McBride SM. 2016. The Phosphotransfer Protein CD1492 Represses Sporulation Initiation in Clostridium difficile. Infect Immun doi:10.1128/IAI.00735-16.

38. Edwards AN, McBride SM. 2017. Determination of the in vitro Sporulation Frequency of Clostridium difficile. Bio Protoc 7.

39. Edwards AN, Nawrocki KL, McBride SM. 2014. Conserved oligopeptide permeases modulate sporulation initiation in Clostridium difficile. Infect Immun 82:4276–91.

40. McBride SM, Sonenshein AL. 2011. Identification of a genetic locus responsible for antimicrobial peptide resistance in Clostridium difficile. Infect Immun 79:167–76.

41. Schmittgen TD, Livak KJ. 2008. Analyzing real-time PCR data by the comparative C(T) method. Nat Protoc 3:1101–8.

42. Edwards AN, Pascual RA, Childress KO, Nawrocki KL, Woods EC, McBride SM. 2015. An alkaline phosphatase reporter for use in Clostridium difficile. Anaerobe 32:98–104.

43. Kirk JA, Fagan RP. 2016. Heat shock increases conjugation efficiency in Clostridium difficile. Anaerobe 42:1–5.

44. Chen X, Katchar K, Goldsmith JD, Nanthakumar N, Cheknis A, Gerding DN, Kelly CP. 2008. A mouse model of Clostridium difficile-associated disease. Gastroenterology 135:1984–92.

45. Pawlowski SW, Calabrese G, Kolling GL, Platts-Mills J, Freire R, AlcantaraWarren C, Liu B, Sartor RB, Guerrant RL. 2010. Murine model of Clostridium difficile infection with aged gnotobiotic C57BL/6 mice and a BI/NAP1 strain. J Infect Dis 202:1708–12.

46. Reeves AE, Koenigsknecht MJ, Bergin IL, Young VB. 2012. Suppression of Clostridium difficile in the gastrointestinal tracts of germfree mice inoculated with a murine isolate from the family Lachnospiraceae. Infect Immun 80:3786–94.

47. Buffie CG, Jarchum I, Equinda M, Lipuma L, Gobourne A, Viale A, Ubeda C, Xavier J, Pamer EG. 2012. Profound alterations of intestinal microbiota following a single dose of clindamycin results in sustained susceptibility to Clostridium difficile-induced colitis. Infect Immun 80:62–73.

48. Theriot CM, Koumpouras CC, Carlson PE, Bergin, II, Aronoff DM, Young VB. 2011. Cefoperazone-treated mice as an experimental platform to assess differential virulence of Clostridium difficile strains. Gut Microbes 2:326–34.

49. Goorhuis A, Bakker D, Corver J, Debast SB, Harmanus C, Notermans DW, Bergwerff AA, Dekker FW, Kuijper EJ. 2008. Emergence of Clostridium difficile infection due to a new hypervirulent strain, polymerase chain reaction ribotype 078. Clin Infect Dis 47:1162–70.

50. Walker AS, Eyre DW, Wyllie DH, Dingle KE, Griffiths D, Shine B, Oakley S, O’Connor L, Finney J, Vaughan A, Crook DW, Wilcox MH, Peto TE, Infections in Oxfordshire Research D. 2013. Relationship between bacterial strain type, host biomarkers, and mortality in Clostridium difficile infection. Clin Infect Dis 56:1589–600.

51. Franke AE, Clewell DB. 1981. Evidence for a chromosome-borne resistance transposon (Tn916) in Streptococcus faecalis that is capable of “conjugal” transfer in the absence of a conjugative plasmid. J Bacteriol 145:494–502.

52. Mullany P, Wilks M, Puckey L, Tabaqchali S. 1994. Gene cloning in Clostridium difficile using Tn916 as a shuttle conjugative transposon. Plasmid 31:320–3.

53. Olling A, Seehase S, Minton NP, Tatge H, Schroter S, Kohlscheen S, Pich A, Just I, Gerhard R. 2012. Release of TcdA and TcdB from Clostridium difficile cdi 630 is not affected by functional inactivation of the tcdE gene. Microb Pathog 52:92–100.

54. El Meouche I, Peltier J, Monot M, Soutourina O, Pestel-Caron M, Dupuy B, Pons JL. 2013. Characterization of the SigD regulon of C. difficile and its positive control of toxin production through the regulation of tcdR. PLoS One 8:e83748.

55. McKee RW, Mangalea MR, Purcell EB, Borchardt EK, Tamayo R. 2013. The second messenger cyclic Di-GMP regulates Clostridium difficile toxin production by controlling expression of sigD. J Bacteriol 195:5174–85.

56. Rocha-Estrada J, Aceves-Diez AE, Guarneros G, de la Torre M. 2010. The RNPP family of quorum-sensing proteins in Gram-positive bacteria. Applied Microbiology and Biotechnology 87:913–923.

57. Cook LC, Federle MJ. 2014. Peptide pheromone signaling in Streptococcus and Enterococcus. Fems Microbiology Reviews 38:473–492.

58. Karlsson S, Burman LG, Akerlund T. 1999. Suppression of toxin production in Clostridium difficile VPI 10463 by amino acids. Microbiology 145 (Pt 7):1683–93.

59. Mani N, Lyras D, Barroso L, Howarth P, Wilkins T, Rood JI, Sonenshein AL, Dupuy B. 2002. Environmental response and autoregulation of Clostridium difficile TxeR, a sigma factor for toxin gene expression. J Bacteriol 184:5971–8.

60. Karlsson S, Dupuy B, Mukherjee K, Norin E, Burman LG, Akerlund T. 2003. Expression of Clostridium difficile toxins A and B and their sigma factor TcdD is controlled by temperature. Infect Immun 71:1784–93.

61. Karlsson S, Burman LG, Akerlund T. 2008. Induction of toxins in Clostridium difficile is associated with dramatic changes of its metabolism. Microbiology 154:3430–6.

62. Bouillaut L, Dubois T, Francis MB, Daou N, Monot M, Sorg JA, Sonenshein AL, Dupuy B. 2019. Role of the global regulator Rex in control of NAD(+) - regeneration in Clostridioides (Clostridium) difficile. Mol Microbiol 111:1671–1688.

63. Martin-Verstraete I, Peltier J, Dupuy B. 2016. The Regulatory Networks That Control Clostridium difficile Toxin Synthesis. Toxins (Basel) 8.

64. Bakker D, Smits WK, Kuijper EJ, Corver J. 2012. TcdC does not significantly repress toxin expression in Clostridium difficile 630DeltaErm. PLoS One 7:e43247.

65. Dupuy B, Sonenshein AL. 1998. Regulated transcription of Clostridium difficile toxin genes. Mol Microbiol 27:107–20.

66. Ransom EM, Kaus GM, Tran PM, Ellermeier CD, Weiss DS. 2018. Multiple factors contribute to bimodal toxin gene expression in Clostridioides (Clostridium) difficile. Mol Microbiol doi:10.1111/mmi.14107.

67. Anjuwon-Foster BR, Tamayo R. 2017. A genetic switch controls the production of flagella and toxins in Clostridium difficile. PLoS Genet 13:e1006701.

68. Hussain HA, Roberts AP, Mullany P. 2005. Generation of an erythromycin-sensitive derivative of Clostridium difficile strain 630 (630Δerm) and demonstration that the conjugative transposon Tn916ΔE enters the genome of this strain at multiple sites. Journal of Medical Microbiology 54:137–141.

69. Thomas CM, Smith CA. 1987. Incompatibility group P plasmids: genetics, evolution, and use in genetic manipulation. Annu Rev Microbiol 41:77–101.

70. Manganelli R, Provvedi R, Berneri C, Oggioni MR, Pozzi G. 1998. Insertion vectors for construction of recombinant conjugative transposons in Bacillus subtilis and Enterococcus faecalis. FEMS Microbiol Lett 168:259–68.

71. McAllister KN, Bouillaut L, Kahn JN, Self WT, Sorg JA. 2017. Using CRISPR- Cas9-mediated genome editing to generate C. difficile mutants defective in selenoproteins synthesis. Sci Rep 7:14672.

